# Temporal composition of the cervicovaginal microbiome associates with hrHPV infection outcomes in a longitudinal study

**DOI:** 10.1101/2023.06.02.543506

**Authors:** Mariano A. Molina, William P. J. Leenders, Martijn A. Huynen, Willem J. G. Melchers, Karolina M. Andralojc

## Abstract

Persistent infections with high-risk human papillomavirus (hrHPV) can cause cervical squamous intraepithelial lesions (SIL) that may progress to cancer. The cervicovaginal microbiome (CVM) correlates with SIL, but the temporal composition of the CVM after hrHPV infections has not been fully clarified. To determine the association between the CVM composition and infection outcome, we applied high-resolution microbiome profiling using the circular probes-based RNA sequencing technology on a longitudinal cohort of cervical smears obtained from 141 hrHPV DNA-positive women with normal cytology at first visit, of whom 51 were diagnosed by cytology with SIL six months later. Here we show that women with a microbial community characterized by low diversity and high *Lactobacillus crispatus* abundance exhibit low risk to SIL development at both visits, while women with a microbial community characterized by high diversity and *Lactobacillus* depletion at first visit have a higher risk of developing SIL. At the level of individual species we observed that an increased abundance for *Gardnerella vaginalis* and *Atopobium vaginae* associate with SIL outcomes at both visits. These species together with *Dialister micraerophilus* showed a moderate discriminatory power for hrHPV infection progression. Our results suggest that the CVM can potentially be used as a biomarker for cervical disease and SIL development after hrHPV infection diagnosis with implications on cervical cancer prevention strategies and treatment of SIL.

## Introduction

High-risk human papillomavirus (hrHPV) infections are associated with premalignant cervical lesions that may progress to cervical cancer (1). Around 80% of all sexually active women will acquire an HPV infection during their lives and in most of the cases the virus is spontaneously cleared by the host immune system (2–4). In some women, however, hrHPV evades the immune response and the infection becomes persistent, promoting the development of squamous intraepithelial lesions (SIL) that eventually can progress to invasive cervical cancer (5, 6). Despite increased use of HPV-vaccines to prevent hrHPV infection, cervical cancer represents a huge public health burden worldwide with over 500,000 diagnoses and over 300,000 deaths yearly (7). Current screening programs include hrHPV DNA testing followed by cytology triage (Pap test). Overall, the clinical specificity of screening is low, resulting in high rates of overdiagnosis and overtreatment, and stratification of women who are at risk of hrHPV-induced cancer remains a challenge (8). Thus, there is a remaining need to better understand the cervicovaginal ecosystem, and to discover and apply effective predictive biomarkers for early detection and treatment of SIL.

The cervicovaginal microbiome (CVM) is a promising candidate biomarker for cervical disease behavior (9, 10). Changes in the composition of the cervicovaginal microbiota have been associated with bacterial vaginosis (BV), pre-term birth, and viral infections caused by HIV and hrHPV (11–15). The CVM is structured in microbial community state types (CSTs) in which specific bacterial species dominate the microbiome or assemble in a diverse microbial population. In a healthy cervix, the CVM is characterized by dominance of *Lactobacillus* species such as *Lactobacillus crispatus* (CST I), while depletion of *Lactobacillus* species and colonization by *Gardnerella vaginalis*, *Atopobium vaginae*, and *Megasphaera genomosp type 1* (CST IV) is typical of dysbiosis (16, 17). CST IV has been associated with hrHPV infection, viral persistence, viral-induced cervical lesions, and cervical cancer (18). In contrast, *Lactobacillus*-dominated microbiomes have been correlated with hrHPV clearance and disease regression (18–20). Most of these observations have been described in cross-sectional studies, and since the CVM is a highly dynamic ecological environment (21), a thorough understanding of how the microbiome changes in the course of hrHPV infection to SIL requires longitudinal microbiome profiling studies.

Evaluating the cervicovaginal microbiota’s role in health and disease mainly relies on 16S rRNA gene sequencing methods (22, 23). Species-level microbiome profiling can be achieved by shotgun metagenomics or circular probes-based RNA sequencing (ciRNAseq) techniques (24–26). The ciRNAseq technology employs single-molecule molecular inversion probes (smMIPs) to target conserved DNA and RNA sequences in the 16S and 18S rRNA genes of microbial species within the CVM. ciRNAseq exhibits high specificity and sensitivity in identifying microbial species in mock community samples and women’s cervical smears (25). Likewise, ciRNAseq provides improved taxonomic resolution compared to 16S rRNA gene sequencing, which is critical for the study of the CVM in hrHPV infections. Through ciRNAseq profiling of the CVM, our group has previously defined associations of the CVM with hrHPV-negative conditions and hrHPV-induced high-grade squamous intraepithelial lesions (HSIL) (25). More recently, we identified subgroups of CSTs based on the abundance of bacterial species commonly overlooked by conventional sequencing methods due to their high level of sequence identity with other species (17, 27, 28). Nonetheless, the temporal associations of these microbial communities and species with hrHPV infection outcomes are unknown.

In this longitudinal study, we investigate the composition of the CVM in a cohort of Dutch women participating in the population-based cervical cancer screening program, with proven hrHPV infection but normal cytology at baseline, who were diagnosed with SIL six-months later or did not develop cervical abnormalities. Our study aimed to evaluate the temporal changes in the microbiome in relation to hrHPV progression in these groups. We show that an initial CST IV-A and high *G. vaginalis* or *A. vaginae* abundance associate with a progressive infection outcome at six-months, while *L. crispatus* dominance at both visits associates with non-progression. In addition to CSTs, we describe a combination of microbial species associated with hrHPV outcomes at both visits and relationships between bacteria occurring in the CVM. Our results suggest that the CVM is a valuable biomarker for hrHPV infection progression.

## Material and methods

### Ethics statement

The Central Committee on Research Involving Human Subjects (CCMO) and the National Institute for Public Health and Environment (RIVM) reviewed and granted approval before the start of the study (No. 2014-1295). All methods were performed in accordance with the Radboudumc ethical guidelines for using human samples, including the Declaration of Helsinki.

### Study subjects and inclusion criteria

A total of 141 women participating in the Dutch population-based cervical cancer screening program and diagnosed with hrHPV infection and cytologically characterized as negative for intraepithelial lesion or malignancy (NILM) were included in the study. Women participating in the screening program were informed that residual material could be used for anonymous research and had the opportunity to opt out. Only residual material from women who did not opt-out was included. At first visit (V1, time = 0 months) and second visit (V2, time = 6 months), 141 cervical smears in PreservCyt were collected, processed, and sequenced for microbiome profiling (25). Five milliliters of each cervical cell suspension were centrifuged for 5 min at 2500×g, and the pellet dissolved in 1 ml of Trizol reagent (Thermo Scientific). RNA was isolated through standard procedures and dissolved in 20 μl nuclease-free water. We routinely processed a maximum of 2 μg of RNA for DNase treatment and cDNA generation, using SuperscriptII (Thermo). From the women with microbiome profiling at V2, a total of 83 cervical smears had sufficient material available for hrHPV DNA testing. The cytological follow-up outcomes at V2 were obtained for all participating women from the nationwide network and registry of histo- and cytopathology in the Netherlands (PALGA; Houten, The Netherlands).

### HrHPV identification and genotyping

DNA from all cervical smears was isolated using MagNA Pure (Roche, Bazel, Switzerland). The purified DNA was eluted in 50 μl TE-buffer (28, 29). HrHPV DNA testing was performed with the Roche Cobas 4800 test, according to the manufacturer’s recommendations in the Department of Medical Microbiology at Radboudumc (30). This test provides separate result for HPV16 and 18, and a pool of 12 other high-risk HPV types (i.e., 31, 33, 35, 39, 45, 51, 52, 56, 58, 59, 66, and 68).

### CiRNAseq microbiome profiling and output analyses

High-resolution microbiome profiling was performed on ∼ 50 ng of cDNA using the ciRNAseq technology (25, 28, 31). Probes (smMIPs) designed to bind to VRs in the 16S and 18S rRNA genes of microbial species in the CVM were mixed with cDNA in a capture hybridization reaction and were circularized via a combined primer extension and ligation reaction. Circularized probes were subjected to PCR with barcoded Illumina primers. After purification of correct-size amplicons, quality control, and quantification (29), a 4 nM library was sequenced on the Illumina Nextseq500 platform (Illumina, San Diego, CA) at the Radboudumc sequencing facility. Reads were mapped against reference regions of interest in our Cervicovaginal Microbiome Panel containing 321 microbial species (25) using the SeqNext module of JSI Sequence Pilot version 4.2.2 build 502 (JSI Medical Systems, Ettenheim, Germany). The settings for reads assigning was a threshold of minimum of 95% of identical bases with the region of interests (ROIs) (25). All identical PCR products were reduced to one consensus read (unique read counts, URC) using a unique molecular identifier. We set an arbitrary threshold of at least 1000 URC from all smMIPs combined in an individual sample, below which we considered an output non-interpretable. For microbial annotation, species with two reactive smMIPs were these both smMIPs had URC. Species with three or more reactive smMIPs were annotated when more than 50% of their specific set of smMIPs had URC (25).

### Microbiome assessment and analyses

Hierarchical clustering (HC) and Partial least-squares discriminant analysis (PLSDA) were performed using ClustVis and MetaboAnalyst, respectively (32, 33). The settings for HC were as follows: clustering distance for columns: Manhattan; clustering method: Ward (34). CSTs designation was performed through unsupervised clustering analyses (28). CSTs were classified into five major groups (I to V) and the subgroups of CSTs I, III, and IV (17, 28) based on microbiome composition.

The predictive diagnostic potential of *A. vaginae*, *G. vaginalis*, *D. micraerophilus*, and *L. crispatus* for distinguishing non-progressive and progressive women at V1 were evaluated by a Random Forest analysis followed by receiver operating characteristic (ROC) curves of the bacterial species markers, and results were quantified by the area under the curve (AUC) using the randomForest (35) and pROC (36) R packages.

SankeyMATIC software was utilized to visualize the temporal changes in microbiomes. Pearson’s r partial correlations between microbial species were determined and generated with the ppcor R package (37). The microbiome variation in the six-month period within a woman was obtained through a Jensen-Shannon distance (JSD) calculation in the philentropy R package (38). JSD values give a measure of similarity between samples (i.e., by calculating the distance between samples) from the same woman. Low JSD values indicate similar microbial communities between samples, and conversely, large values indicate less similar communities.

### Statistical analysis

GraphPad Prism v9.4.0 (GraphPad Software, Inc., USA) was used to analyze datasets and determine the Shannon’s diversity indices and Odd ratios. The statistical significance of differences was calculated using the Kruskal-Wallis test for multiple comparisons followed by a Benjamini-Hochberg test correction. Mann-Whitney U and Wilcoxon rank tests were employed for single and paired analyses, respectively. A McNemar’s test with a continuity correction was applied for matched-pairs analyses between both visits.

## Results

### Study design and hrHPV infection outcomes

Cervical smears from 141 women with DNA confirmed hrHPV infection and a cytological diagnosis of negative for intraepithelial lesion or malignancy (NILM) were profiled for CVM at first visit (V1). Of these, 90 women also had a diagnosis NILM at 6 months (63.8%) (non-progression group, NP) while 51 women (36.2%) were diagnosed with low-grade squamous intraepithelial lesions (LSIL) (41/51, 80.4%) or HSIL (10/51, 19.6%) (progression group, P) (Figure 1).

**Figure 1.**
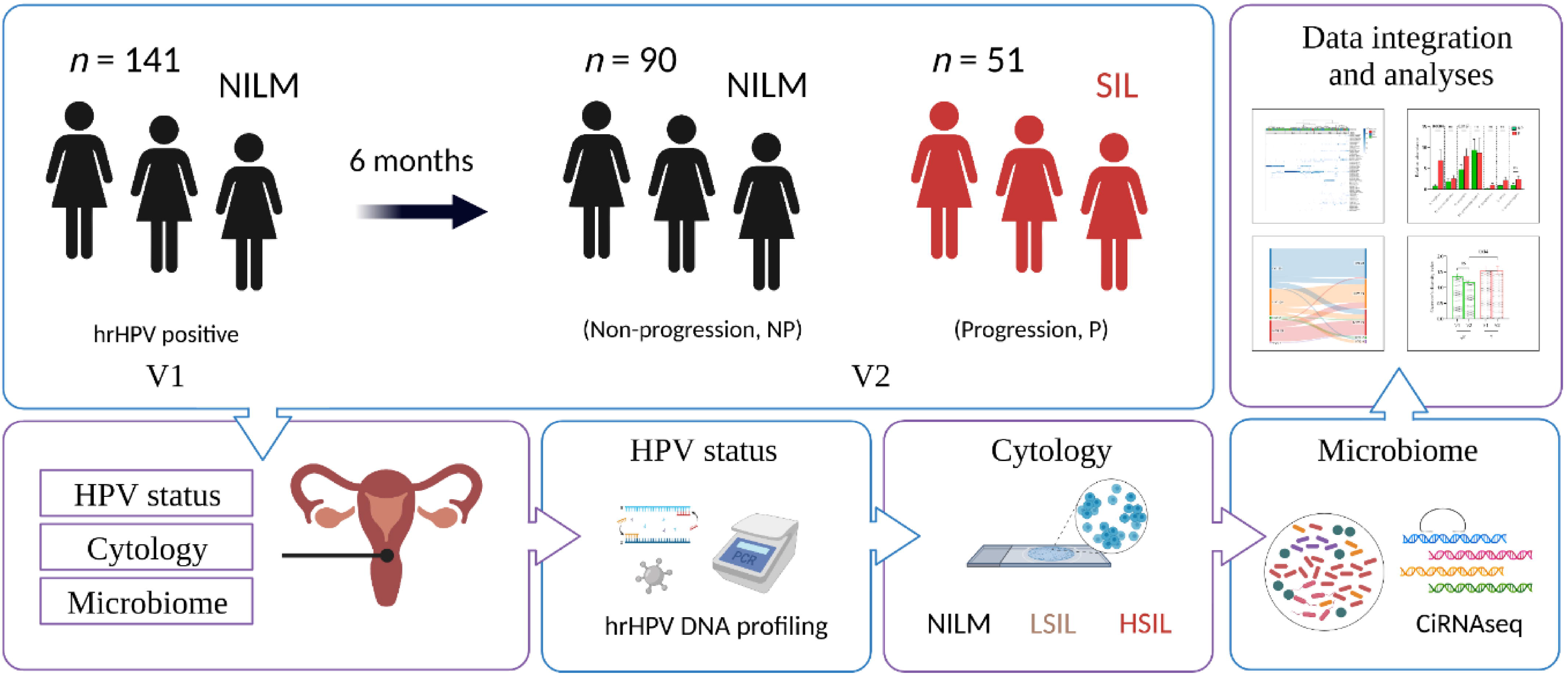
Study design. All 141 women entered the study at baseline with DNA confirmed hrHPV infection, no cervical abnormalities and CVM profiling. The results of follow-up cytology were assessed at 6 months to determine whether the individual had progressed to intraepithelial lesion or malignancy (LSIL, HSIL) or not (NILM). By 6 months, 90 women were confirmed for NILM, and 52 women had a LSIL (n = 40) or HSIL (n = 10) diagnosis. Experimental procedures, analysis, and integration were carried out as described in Material and methods.

### Early microbiome composition and hrHPV infection outcomes

Through unsupervised cluster analysis, we characterized the composition of the CVM in our longitudinal cohort at baseline (*n* = 141) and determined their association with cytological outcomes at six-months. Microbiomes clustered in CSTs dominated by *Lactobacillus* species: clusters I, III, and V (Figure 2a, left clusters), and CSTs with a high diversity: clusters II and IV (Figure 2a, right clusters, and Supplementary Figure 1a), including the subgroups of CSTs I (I-A, I-B), III (III-A, III-B) and IV (IV-A, IV-B, and IV-C) (28). We did not observe a significant association between the overall baseline *Lactobacillus*-dominated (LDO, CSTs I, II, III, and V combined) and *Lactobacillus*-depleted (LDE, CSTs IV combined) microbiomes with hrHPV infection outcomes at V2 (Figures 2a-2b). Nevertheless, we see a clear trend where, of the CST types, CST I-A at baseline was most strongly associated with NILM at six-months (26/32, 81.2%, OR 0.32, 95% CI 0.12–0.82, *p* = 0.03, *q* = 0.15, Fisher’s exact test), while CST IV-A at baseline was most strongly associated with SIL outcomes at six-months (9/15, 60%, OR 3.07, 95% CI 1.03–9.40, *p* = 0.04, *q* = 0.16), however, the associations were only moderate when corrected for multiple testing (FDR < 0.2) (Figures 2a-2b).

**Figure 2.**
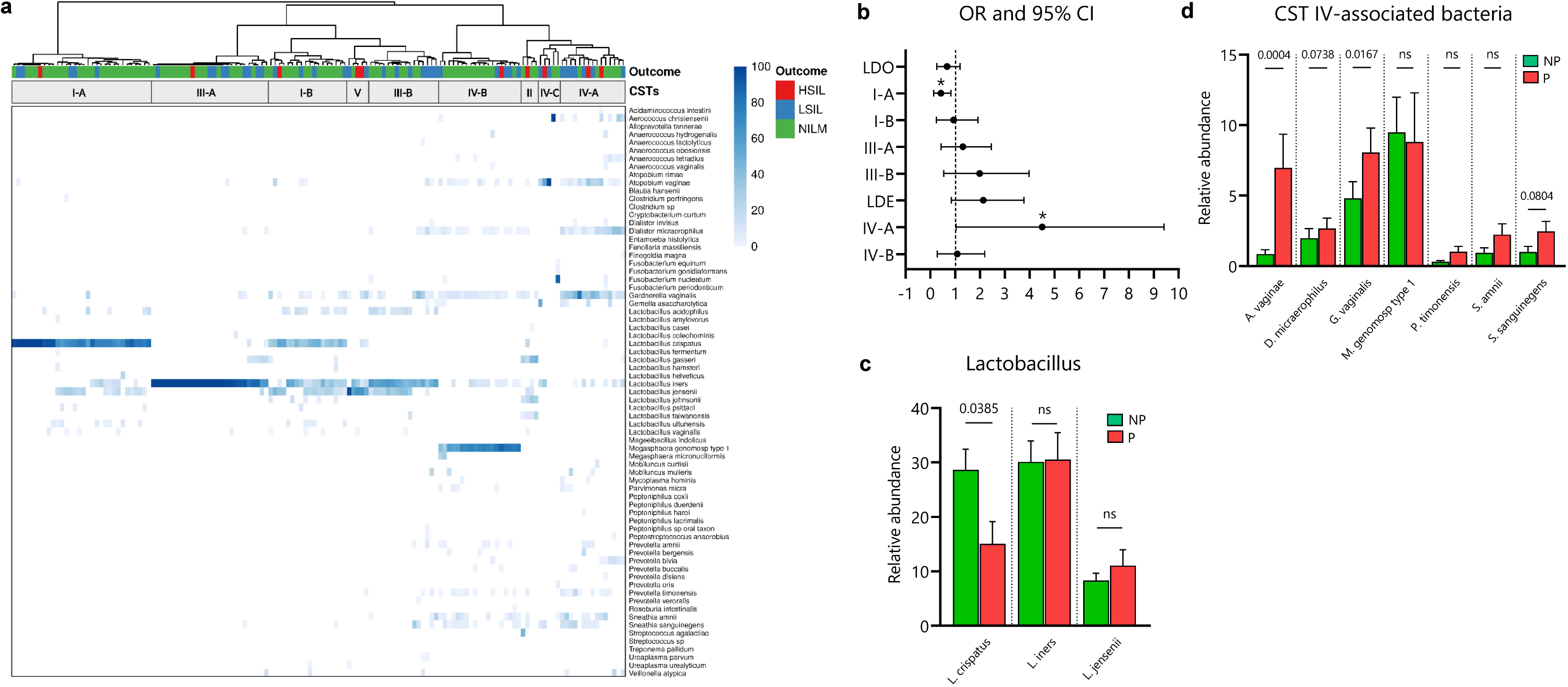
Early cervicovaginal microbiome composition associates with hrHPV infection outcomes at six-months. **a.** Cluster analysis of species-level profiling of the cervicovaginal microbiota at first collection visit (V1). Visualization of the distribution of hrHPV infection outcomes based on clusters show enrichment of NILM and SIL outcomes in specific communities. **b.** Odd ratios (OR) and 95% confidence intervals comparing baseline CST groups (LDO: I, II, III, and V; LDE: IV) and individual CST subgroups for hrHPV infection progression at six-months. Analyses of the relative abundances of the most abundant bacterial species in the CVM at V1 demonstrate association of *Lactobacillus crispatus* with non-progression **(c)** and pathogenic anaerobes with the infection outcomes at six-months **(d)**. OR in b were analyzed through a Fisher’s exact test, * *p* < 0.05. Differences in relative abundances were analyzed by using a Kruskal-Wallis test followed by the Benjamini-Hochberg test correction for multiple comparisons. *q* values are shown in **c** and **d** and error bars represent standard error of the mean ± s.e.m. NP = non-progression group; P = progression group; LDO = *Lactobacillus*-dominated; LDE = *Lactobacillus*-depleted.

To further explore the association of the CVM with progression to SIL, we examined it at level of individual bacterial species. We observed a significantly increased abundance of *L. crispatus* (*q* = 0.03, Kruskal Wallis test) in the NP group when compared to the P group (Figure 2c). Moreover, we noticed that *A. vaginae* (*q* = 0.0004, Kruskal Wallis test), *G. vaginalis* (*q* = 0.01), *D. micraerophilus* (*q* = 0.07) and S*. sanguinegens* (*q* = 0.08), which are typical species found in CST IV, were more abundant in the P group than in the NP group (Figure 2d).

### Dynamics of the microbiome and hrHPV infection outcomes

Six months after the initial diagnosis of hrHPV infection (V2), we again performed microbiome profiling on the cervical smears from all participating women (*n* = 141). We then established their CVM composition and examined the microbial changes between both visits and their association with hrHPV infection outcomes (Supplementary Figure 2). Although we did not observe a significant association between CST subgroups and hrHPV infection outcomes at V2 like we observed at V1 (Figure 2, Supplementary Figure 2), we found that, LDE CSTs (IV) correlated with SIL (OR 2.21, 95% CI 1.02–4.45, *p* = 0.03, Fisher’s exact test), while LDO (I, II, III, and V) were associated with NILM (OR 0.45, 95% CI 0.22–0.97) (Figure 3b).

**Figure 3.**
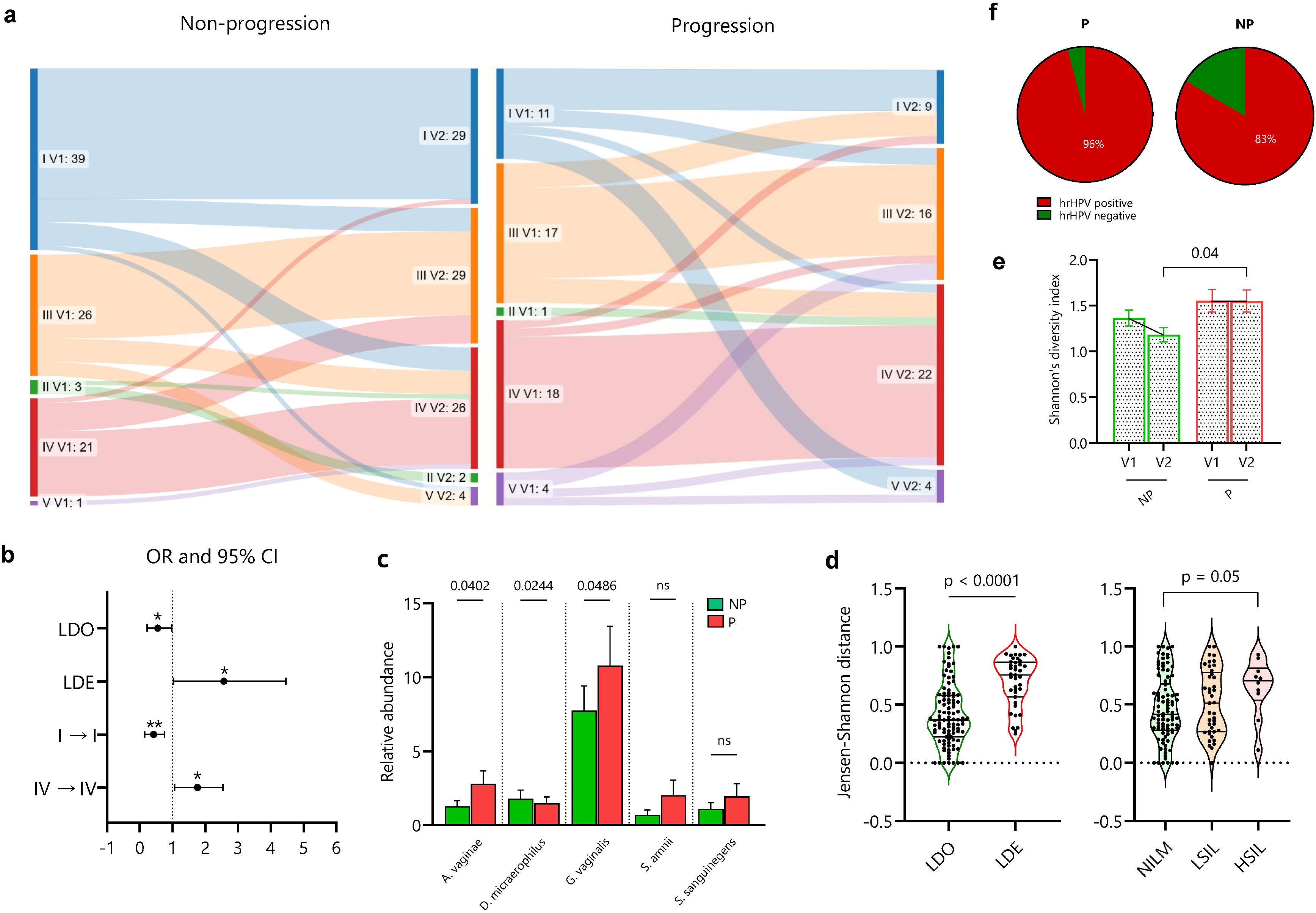
Dynamics of the microbiome and hrHPV infection outcomes. **a.** The microbial shifts between both visits and groups. **b.** Odd ratios (OR) and 95% confidence intervals comparing CST groups (LDO: I, II, III, and V; LDE: IV) at V2 and the six-months stability of CSTs I and IV with hrHPV infection outcomes. **c.** Similarity of the CVM composition per microbial community and infection outcome through the Jensen-Shannon distance (JSD). **d.** Comparison of relative abundances of the most abundant bacterial species associated with LDE microbiomes between NP and P groups. **e.** Comparison of Shannon’s diversity indices for all microbiomes in NP and P groups at both visits. **f.** Analysis on the hrHPV status in a subcohort of 83 women with hrHPV DNA testing at V2. OR in **b** were analyzed through a Fisher’s exact test, * *p* < 0.05, ** *p* < 0.01. Differences in relative abundances, JSD per outcomes, and Shannon indices per group, were analyzed by using a Kruskal-Wallis test followed by the Benjamini-Hochberg test correction for multiple comparisons. *q* values are shown in **d** and **e** and error bars represent standard error of the mean ± s.e.m. Differences in JSD by Lactobacillus composition and paired Shannon indices were analyzed by using a Mann–Whitney U test and Wilcoxon matched-pairs test, respectively. NP = non-progression group; P = progression group; LDO = *Lactobacillus*-dominated; LDE = *Lactobacillus*-depleted.

We did not find a significant association between the CSTs II, III, IV, and V with microbiome stability in both groups (Figure 3a). Nonetheless, compared to these CSTs in the NP group, CST I was significantly associated with a stable composition between both visits (*p* = 0.01, McNemar’s test) (Figure 3a). Moreover, when analyzing the association between the six-months stability of CSTs with the hrHPV outcomes, we found that compared to other CSTs, stable CST I (OR 0.24, 95% CI 0.09–0.64, *p* = 0.003, Fisher’s exact test) and CST IV (OR 2.48, 95% CI 1.12–5.66, *p* = 0.03) were significantly associated with non-progression and progression, respectively (Figure 3b).

Next, we examined the association of the CVM composition at V2 with the infection outcomes by comparing the relative abundances of pathogenic anaerobic species between both groups. Interestingly, we observed that the species *D. micraerophilus* (*q* = 0.02, Kruskal Wallis test) was more abundant in women with NILM than in women with SIL (Figure 3c). Alternatively, we found that *A. vaginae* (*q* = 0.04), *G. vaginalis* (*q* = 0.04), *P. bivia* (*q* = 0.008) and *P. buccalis* (*q* = 0.008) were more abundant in the SIL group than in the NILM group (Figures 3c and Supplementary Figure 3).

To further evaluate the association of the temporal CVM composition with hrHPV infection outcomes, we calculated the Jensen-Shannon distance (JSD) (38) of the microbiome composition between both visits. A low JSD indicates a high similarity in microbial composition between the timepoints. We observed that women with LDO at baseline had a significantly more similar microbiome at V2 than women with LDE at baseline (*p* <0.0001, Mann–Whitney U test) (Figure 3d). Conversely, women with NILM outcomes had a more similar microbiome composition between both visits than those who developed HSIL (*p* = 0.05, *q* = 0.17, Kruskal Wallis test) (Figure 3d).

We then assessed the Shannon’s diversity index and observed that women in both NP and P groups did not exhibit a significant change in microbial diversity from V1 to V2 (Wilcoxon matched-pairs test) (Figure 3e). Nevertheless, women in the NP group had a significantly lower microbial diversity than the P group at V2 (*q* = 0.04, Kruskal Wallis test) (Figure 3e). To test whether these microbial dynamics were associated with hrHPV persistence, we analyzed the hrHPV status for a subcohort of cervical smears with available hrHPV DNA testing at V2 (*n* = 83) and noticed that there were more hrHPV positive cervical smears in women with SIL (23/24, 96%) than women with NILM (49/59, 83%) (OR 4.69, 95% CI 0.68– 53.04, *p* = 0.16, Fisher’s exact test) (Figure 3f). Altogether, these findings demonstrate that prolonged *Lactobacillus* depletion, high microbial diversity, and increased abundance of CST IV-associated bacteria correlate with SIL. Conversely, a stable microbiome composition characterized by *Lactobacillus* dominance and low microbial diversity over a six-months period correlates with non-progression.

### Microbiome-based prediction of hrHPV infection outcomes

Aside from estimating species that most significantly associate with the NP and P groups, we aimed to determine to what extend a combination of species associated with each group at both visits and whether such combination could be used to predict hrHPV infection outcomes at V1. To this purpose, we first performed a Partial least-squares discriminant analysis (PLSDA) with all microbiomes collected at V1 and V2 (*n* = 141) (Figure 4a). We determined that *L. crispatus*, *A. vaginae*, *D. micraerophilus*, and *G. vaginalis* showed the strongest correlations with PLSDA Component 1 at both visits. Analyses of the Variable Importance in Projection (VIP) scores, a weighted sum of squares of the PLS loadings, showed consistent microbial species separating both groups in PLSDA C1 and the relative abundance associated with each group (Supplementary Figure 4).

**Figure 4.**
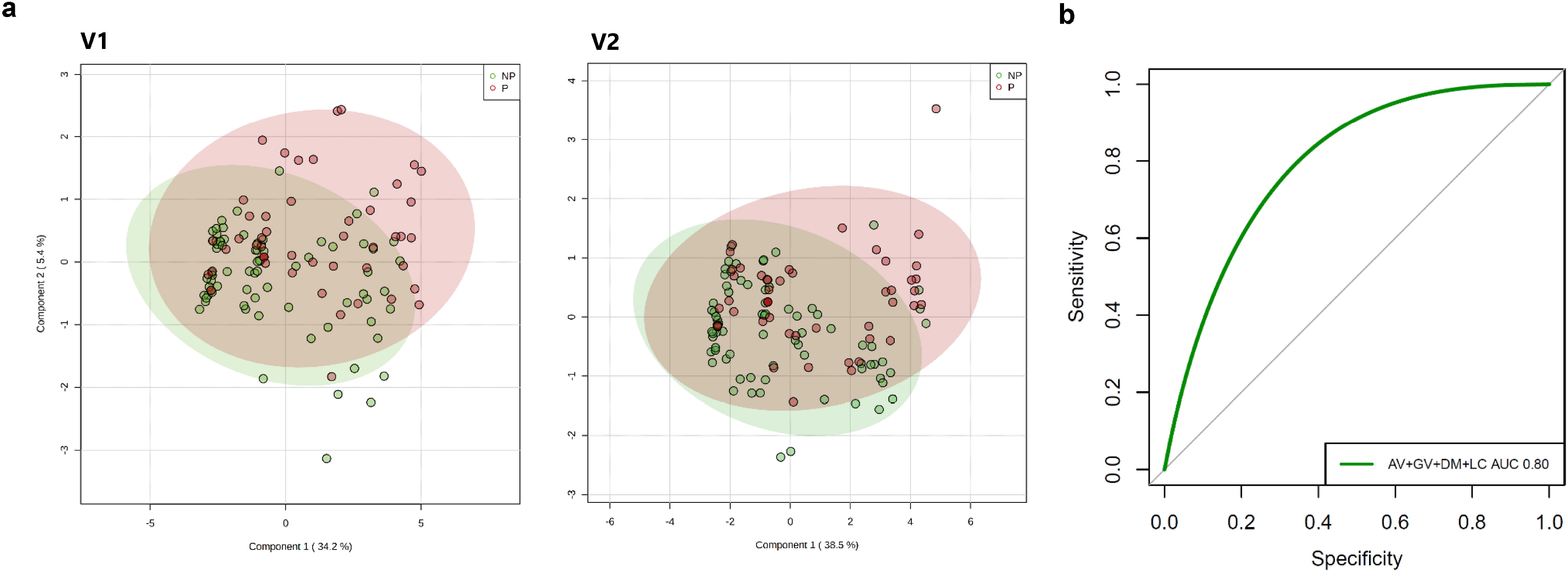
Cervicovaginal microbial species associated with hrHPV infection outcomes. **a.** Partial least-squares discriminant analysis (PLSDA) of women’s CVM (*n* = 141) shows a similar separation between NP and P groups at both V1 and V2. **b.** Receiver operating characteristics (ROC) curve and AUC of *A. vaginae* (AV), *G. vaginalis* (GV), *D. micraerophilus* (DM), and *L. crispatus* (LC) abundances in the microbiome from 141 hrHPV-positive women at V1 were calculated to generate a predictive model for the infection outcomes at six-months.

Next, we performed a Random Forest analysis and ROC to assess the performance of the early abundances in the CVM of the species *L. crispatus*, *A. vaginae*, *D. micraerophilus*, and *G. vaginalis*, which exhibited the strongest associations in our PLSDA (Figure 4a), in predicting hrHPV outcomes at six-months. We found that these species together had a moderate discriminatory power at baseline for hrHPV infection progression at V2 with an AUC of 0.80 (95% CI 0.64–0.96) (Figure 4b).

### Correlations among the cervicovaginal microbiota in hrHPV infections

Lastly, to assess whether relationships between bacterial species were observed and persisted in non-progressive and progressive microbiomes, we performed Pearson’s partial r correlation analyses considering the most abundant species in both NP and P groups. We analyzed the microbial species abundances by integrating the two timepoints datasets to establish the correlations that persisted throughout both visits. In the NP group, there was a positive correlation between *A. vaginae* and *D. micraerophilus* and *G. vaginalis*. In the P group, there were inverse relationships between *Lactobacillus* and *A. vaginae*, between *M. genomosp type 1* and other pathogenic bacteria, and between *Prevotella* species (Figure 5). Alternatively, we observed significant positive correlations between *D. micraerophilus* and *Prevotella* species and between *Prevotella* and *Sneathia* species in the P group (Figure 5). In both groups, there was a negative correlation between *Lactobacillus* and *G. vaginalis* and *M. genomosp type 1* (Figure 5). Likewise, both groups exhibited a significantly positive correlation between *D. micraerophilus* and *P. bivia* (Figure 5). In conclusion, associations between *Lactobacillus* and pathogenic anaerobes, and between pathogenic anaerobes themselves, appear typical in the CVM during hrHPV infections, and we observe no indication that the species-species associations are typical for either the P or the NP group.

**Figure 5.**
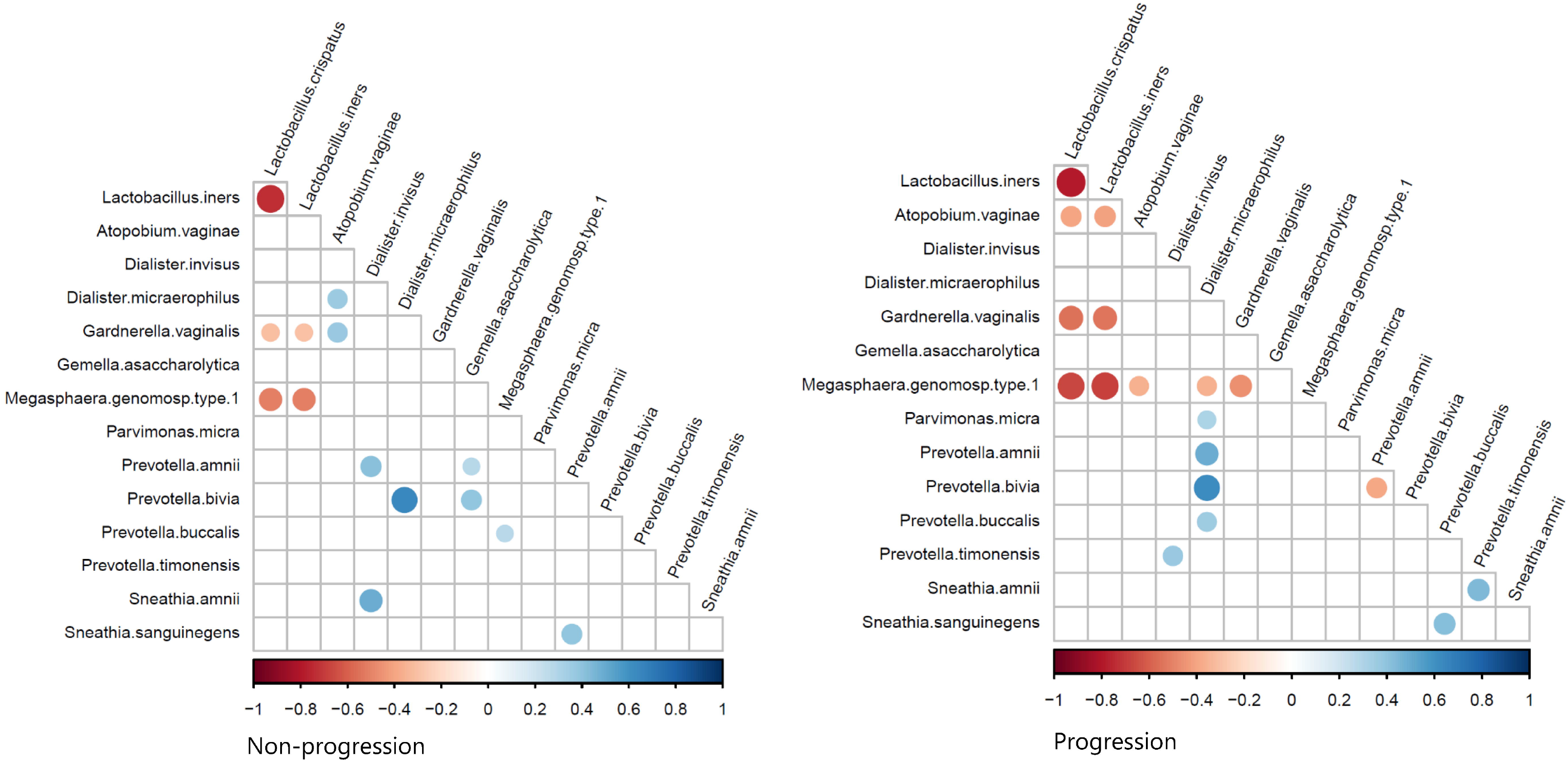
Interdependent relationships between the cervicovaginal microbiota in hrHPV infection. Pearson’s r partial correlations for multiple comparisons were estimated with the most abundant bacterial species in the microbiome of women in NP and P groups and integrating V1 and V2 datasets (*n* = 141). Color, size, and shade indicate the extent of positive and negative correlations. Correlation significance: *p* < 0.001

## Discussion

The composition of the CVM not only correlates with hrHPV infections and cervical disease, but it may also predict the infection outcome. In this study, we observed that following hrHPV infection diagnosis, women with the microbial community IV-A (17, 28), characterized by *G. vaginalis* dominance and co-occurrence with *D. micraerophilus*, *A. vaginae*, *S. amnii*, and *S. sanguinegens* associate with progression to SIL six-months later. Pathogenic anaerobes such as *G. vaginalis* have been associated with viral persistence and cervical lesions (17, 19, 39). Our findings are in line with previous longitudinal studies that established an association between bacterial vaginosis (BV) and the species *G. vaginalis* with cervical neoplastic lesions (19, 40). Notably, we found that *A. vaginae* abundance at both visits is associated with infection progression and SIL at second visit. Increased *A. vaginae* abundance in the microbiome has been reported as a hallmark of SIL development (41–43). *A. vaginae* induces cytotoxic immune responses in cervicovaginal epithelial cells that reduces the protective mucosal layer, which might facilitate hrHPV persistence and integration into host cells (44, 45). Thus, since these bacteria are highly abundant in hrHPV progressive infections and SIL, they could potentially be applied as biomarkers for cervical carcinogenesis. Moreover, these pathogenic species may represent promising targets for microbiome-based therapies against the development of cervical cancer (17).

Microbiomes abundant in *Lactobacillus* are associated with cervical health and their depletion results in cervical disorders (10, 13, 41, 46, 47). *Lactobacillus* species create a favorable microenvironment that allows sustained presence of lactate-producing bacteria and prevent outgrowth of harmful bacteria such as *G. vaginalis* (17, 48, 49). By this mechanism, *Lactobacillus* species may therefore prevent dysbiosis and persistent hrHPV infections (17, 50). Additionally, microbial dominance by *L. crispatus* (CST I), *L. gasseri* (CST II), or *L. jensenii* (CST V) has been associated with hrHPV negative conditions, viral clearance, and regression of cervical lesions (17, 20). Although we did not assess this relationship in CSTs II and V due to the low prevalence of these CSTs in this study, these *Lactobacillus* species are known to stimulate a non-inflammatory state in the cervical epithelium, which facilitates effective immune responses against hrHPV infections and carcinogenesis (17, 51). Similarly, we described that LDO microbiomes have a more stable composition than LDE microbiomes. Since non-progressive women have microbial communities richer in *Lactobacillus* species than progressive women, this may explain the protection against hrHPV progression observed in our study (21, 52).

Aside from disease development, the microbiome dynamics also rely in the interactions of the cervicovaginal microbiota with the virus and the microenvironment (40, 53). In our study, we describe that *Lactobacillus* species exhibit a strong negative relationship with CST IV-associated bacteria, which can be explained by the ecological conditions where they grow and their antimicrobial activities against pathogenic bacteria (17, 54, 55). Interestingly, *D. micraerophilus* showed strong positive associations with several *Prevotella* species in women with SIL outcomes, and in particular, a strong relationship with *P. bivia* was observed in both NP and P groups. *Prevotella* species often co-occur with *D. micraerophilus* and other pathogenic anaerobes in the CVM, and it is a clear example of how microbial species can associate with each other synergistically in the cervicovaginal environment (17, 56). *P. bivia* is an important source of ammonia and sialidase in the vaginal mucus and has been associated with cervical disease, which may explain its occurrence during infection (56). Similarly, we observed that *P. timonensis* positively correlated with *S. amnii* in women with SIL outcomes (Figure 5). *P. timonensis* interacts with vaginal dendritic cells, which are involved in mucosal inflammation (57, 58), and both species have been associated with viral persistence, slower regression of SIL, and cervical cancer (20, 59, 60). *Sneathia* and *Prevotella* species also express several homologous genes that are enriched in CST IV and that allow them to consume glycogen and mucins from cervical cells (61). It could be hypothesized that *S. amnii* or *P. timonensis* may facilitate each other’s colonization, contributing to the risk of neoplastic lesions in hrHPV infection. This hypothesis is also consistent with the hrHPV downregulation of immune peptides that act as amino acid sources for *Lactobacillus* species, which leads to the growth of CST IV-associated bacteria in the CVM (62). In general, the microbial dynamics during hrHPV infections remain interesting markers for infection behavior, but further studies are clearly needed to validate these associations *in vitro* and *in vivo*.

The strengths of the study are the use of the ciRNAseq technology for targeted sequencing of the microbiome and the application of longitudinal profiling in our study cohort. Potential limitations may include a short study period (six months) and a relatively small cohort size. Of note, although we included women with LSIL outcomes in our progression group, LSIL are considered non-progressive lesions (63), and therefore the microbiome associations described here should be considered carefully when investigating outcomes beyond six-months. Women with LSIL, however, developed these cervical abnormalities at V2 from diagnosed NILM at V1, which is defined as hrHPV infection progression. We were also unable to control for phase of the menstrual cycle or antibiotic use during the study, which may impact on CST composition (64). Therefore, a larger cohort and longitudinal clinical studies will be needed to validate our findings.

In summary, we have shown how bacterial species, communities, dynamics, and relationships are relevant for assessing the role of the CVM in hrHPV carcinogenesis. This way, the CVM could be employed to support current cervical cancer prevention strategies and therapies against cervical lesions. Nonetheless, even though CSTs correlate with the infection outcome, their usefulness as biomarkers for cervical disease is not clear yet. Our in-depth analyses suggest species like *A. vaginae*, *G. vaginalis*, *L. crispatus*, and *D. micraerophilus* exhibiting strong associations with cervical conditions, and clustering species into CSTs does not necessarily result in better biomarkers than just examining the presence of a few species (65). Further supervised analyses like Random Forest integrating host cell gene expression (66) to the microbiome data would be valuable to obtain a combination of biomarkers for disease progression, and such studies are on the way. Since ciRNAseq can provide bacterial information on DNA and RNA levels (25), while simultaneously perform transcriptome profiling (66), it could be applied to better understand the relationship of the microbiome with hrHPV infections.

## Acknowledgements

Availability of data.

The sequence read data generated in this study are available at NCBI in the Sequencing Read Archive, projects PRJNA856437 (174 files) (67) and PRJEB45937 (108 files) (68), Sample Accession Numbers are shown in Supplementary File 3).

## Authorship and authors’ contributions

All authors have read the journal’s authorship agreement. KA, WM, and WL conceptualized the study. MM performed the data analyses for the manuscript, supervised by KA, MH, WM, and WL. MM drafted the manuscript and was revised by all authors (KA, MH, WL, and WM). All authors approved the manuscript and contributed to the final version for publication.

BioRender.com was used to design figures for the manuscript.

## Abbreviations

AUC: area under the curve
BV: bacterial vaginosis
CiRNAseq: circular probes-based RNA sequencing
CSTs: community state types
CVM: cervicovaginal microbiome
HC: Hierarchical clustering
hrHPV: high-risk human papillomavirus
HSIL: high-grade squamous intraepithelial lesions
JSD: Jensen-Shannon distance
LDO: Lactobacillus-dominated
LDE: Lactobacillus-depleted
LSIL: low-grade squamous intraepithelial lesions
NILM: negative for intraepithelial lesions or malignancy
NP: non-progression
P: progression
PLSDA: Partial least squares discriminant analysis
ROC: receiver operating characteristic
SIL: squamous intraepithelial lesions
URC: unique read counts

## References

1. Hausen Hz. Papillomaviruses Causing Cancer: Evasion From Host-Cell Control in Early Events in Carcinogenesis. JNCI: Journal of the National Cancer Institute. 2000;92(9):690–8.

2. Doorbar J, Quint W, Banks L, Bravo IG, Stoler M, Broker TR, et al. The biology and life-cycle of human papillomaviruses. Vaccine. 2012;30 Suppl 5:F55–70.

3. Stanley M. Immune responses to human papillomavirus. Vaccine. 2006;24:S16–S22.

4. Sasagawa T, Takagi H, Makinoda S. Immune responses against human papillomavirus (HPV) infection and evasion of host defense in cervical cancer. Journal of Infection and Chemotherapy. 2012;18(6):807–15.

5. Steinbach A, Riemer AB. Immune evasion mechanisms of human papillomavirus: An update. International Journal of Cancer. 2018;142(2):224–9.

6. Koshiol J, Lindsay L, Pimenta JM, Poole C, Jenkins D, Smith JS. Persistent human papillomavirus infection and cervical neoplasia: a systematic review and meta-analysis. Am J Epidemiol. 2008;168(2):123–37.

7. Molina MA, Carosi Diatricch L, Castany Quintana M, Melchers WJG, Andralojc KM. Cervical cancer risk profiling: molecular biomarkers predicting the outcome of hrHPV infection. Expert Rev Mol Diagn. 2020:1–22.

8. Wang R, Pan W, Jin L, Huang W, Li Y, Wu D, et al. Human papillomavirus vaccine against cervical cancer: Opportunity and challenge. Cancer Letters. 2020;471:88–102.

9. Curty G, de Carvalho PS, Soares MA. The Role of the Cervicovaginal Microbiome on the Genesis and as a Biomarker of Premalignant Cervical Intraepithelial Neoplasia and Invasive Cervical Cancer. International journal of molecular sciences. 2019;21(1):222.

10. Ventolini G, Vieira-Baptista P, De Seta F, Verstraelen H, Lonnee-Hoffmann R, Lev-Sagie A. The Vaginal Microbiome: IV. The Role of Vaginal Microbiome in Reproduction and in Gynecologic Cancers. J Low Genit Tract Dis. 2022;26(1):93–8.

11. Chen X, Lu Y, Chen T, Li R. The Female Vaginal Microbiome in Health and Bacterial Vaginosis. Front Cell Infect Microbiol. 2021;11:631972-.

12. Feehily C, Crosby D, Walsh CJ, Lawton EM, Higgins S, McAuliffe FM, et al. Shotgun sequencing of the vaginal microbiome reveals both a species and functional potential signature of preterm birth. npj Biofilms and Microbiomes. 2020;6(1):50.

13. Mitra A, MacIntyre DA, Lee YS, Smith A, Marchesi JR, Lehne B, et al. Cervical intraepithelial neoplasia disease progression is associated with increased vaginal microbiome diversity. Sci Rep. 2015;5(1):16865.

14. Petrova MI, van den Broek M, Balzarini J, Vanderleyden J, Lebeer S. Vaginal microbiota and its role in HIV transmission and infection. FEMS Microbiology Reviews. 2013;37(5):762–92.

15. Kyrgiou M, Mitra A, Moscicki A-B. Does the vaginal microbiota play a role in the development of cervical cancer? Translational Research. 2017;179:168–82.

16. Ravel J, Gajer P, Abdo Z, Schneider GM, Koenig SSK, McCulle SL, et al. Vaginal microbiome of reproductive-age women. Proc Natl Acad Sci. 2011;108(Supplement 1):4680.

17. Molina Mariano A, Coenen Britt A, Leenders William PJ, Andralojc Karolina M, Huynen Martijn A, Melchers Willem JG. Assessing the Cervicovaginal Microbiota in the Context of hrHPV Infections: Temporal Dynamics and Therapeutic Strategies. mBio. 2022;0(0):e01619–22.

18. Norenhag J, Du J, Olovsson M, Verstraelen H, Engstrand L, Brusselaers N. The vaginal microbiota, human papillomavirus and cervical dysplasia: a systematic review and network meta-analysis. BJOG: An International Journal of Obstetrics & Gynaecology. 2020;127(2):171–80.

19. Brotman RM, Shardell MD, Gajer P, Tracy JK, Zenilman JM, Ravel J, et al. Interplay between the temporal dynamics of the vaginal microbiota and human papillomavirus detection. J Infect Dis. 2014;210(11):1723–33.

20. Mitra A, MacIntyre DA, Ntritsos G, Smith A, Tsilidis KK, Marchesi JR, et al. The vaginal microbiota associates with the regression of untreated cervical intraepithelial neoplasia 2 lesions. Nature Communications. 2020;11(1):1999.

21. Ravel J, Brotman RM, Gajer P, Ma B, Nandy M, Fadrosh DW, et al. Daily temporal dynamics of vaginal microbiota before, during and after episodes of bacterial vaginosis. Microbiome. 2013;1(1):29.

22. Clarridge JE. Impact of 16s rRNA gene sequence analysis for identification of bacteria on clinical microbiology and infectious diseases. Clin Microbiol Rev. 2004;17(4):840.

23. Graspeuntner S, Loeper N, Künzel S, Baines JF, Rupp J. Selection of validated hypervariable regions is crucial in 16S-based microbiota studies of the female genital tract. Sci Rep. 2018;8(1):9678.

24. Yang Q, Wang Y, Wei X, Zhu J, Wang X, Xie X, et al. The alterations of vaginal microbiome in hpv16 infection as identified by shotgun metagenomic sequencing. Front Cell Infect Microbiol. 2020;10(286).

25. Andralojc KM, Molina MA, Qiu M, Spruijtenburg B, Rasing M, Pater B, et al. Novel high-resolution targeted sequencing of the cervicovaginal microbiome. BMC Biology. 2021;19(1):267.

26. Noecker C, McNally CP, Eng A, Borenstein E. High-resolution characterization of the human microbiome. Translational Research. 2017;179:7–23.

27. Kullen MJ, Sanozky-Dawes RB, Crowell DC, Klaenhammer TR. Use of the DNA sequence of variable regions of the 16S rRNA gene for rapid and accurate identification of bacteria in the Lactobacillus acidophilus complex. Journal of Applied Microbiology. 2000;89(3):511–6.

28. Molina MA, Andralojc KM, Huynen MA, Leenders WPJ, Melchers WJG. In-depth insights into cervicovaginal microbial communities and hrHPV infections using high-resolution microbiome profiling. npj Biofilms and Microbiomes. 2022;8(1):75.

29. van den Heuvel CNAM, Loopik DL, Ebisch RMF, Elmelik D, Andralojc KM, Huynen M, et al. RNA-based high-risk HPV genotyping and identification of high-risk HPV transcriptional activity in cervical tissues. Modern Pathology. 2020;33(4):748–57.

30. Heideman DAM, Hesselink AT, Berkhof J, van Kemenade F, Melchers WJG, Daalmeijer NF, et al. Clinical Validation of the cobas 4800 HPV Test for Cervical Screening Purposes. Journal of Clinical Microbiology. 2011;49(11):3983–5.

31. de Bitter T, van de Water C, van den Heuvel C, Zeelen C, Eijkelenboom A, Tops B, et al. Profiling of the metabolic transcriptome via single molecule molecular inversion probes. Sci Rep. 2017;7(1):11402.

32. Metsalu T, Vilo J. ClustVis: a web tool for visualizing clustering of multivariate data using Principal Component Analysis and heatmap. Nucleic Acids Res. 2015;43(W1):W566–W70.

33. Xia J, Psychogios N, Young N, Wishart DS. MetaboAnalyst: a web server for metabolomic data analysis and interpretation. Nucleic Acids Research. 2009;37(suppl_2):W652-W60.

34. Jiang X, Hu X, He T. Identification of the clustering structure in microbiome data by density clustering on the Manhattan distance. Science China Information Sciences. 2016;59(7):070104.

35. Breiman L. Random Forests. Machine Learning. 2001;45(1):5–32.

36. Robin X, Turck N, Hainard A, Tiberti N, Lisacek F, Sanchez J-C, et al. pROC: an open-source package for R and S+ to analyze and compare ROC curves. BMC Bioinformatics. 2011;12(1):77.

37. Kim S. ppcor: An R Package for a Fast Calculation to Semi-partial Correlation Coefficients. CSAM. 2015;22(6):665–74.

38. Drost H-G. Philentropy: Information Theory and Distance Quantification with R. Journal of Open Source Software. 2018;3(26).

39. Di Paola M, Sani C, Clemente AM, Iossa A, Perissi E, Castronovo G, et al. Characterization of cervico-vaginal microbiota in women developing persistent high-risk Human Papillomavirus infection. Scientific Reports. 2017;7(1):10200.

40. Arokiyaraj S, Seo SS, Kwon M, Lee JK, Kim MK. Association of cervical microbial community with persistence, clearance and negativity of Human Papillomavirus in Korean women: a longitudinal study. Scientific Reports. 2018;8(1):15479.

41. So KA, Yang EJ, Kim NR, Hong SR, Lee J-H, Hwang C-S, et al. Changes of vaginal microbiota during cervical carcinogenesis in women with human papillomavirus infection. PLoS One. 2020;15(9):e0238705.

42. Seo S-S, Oh HY, Lee J-K, Kong J-S, Lee DO, Kim MK. Combined effect of diet and cervical microbiome on the risk of cervical intraepithelial neoplasia. Clinical Nutrition. 2016;35(6):1434–41.

43. Zhou F-Y, Zhou Q, Zhu Z-Y, Hua K-Q, Chen L-M, Ding J-X. Types and viral load of human papillomavirus, and vaginal microbiota in vaginal intraepithelial neoplasia: a cross-sectional study. Annals of Translational Medicine. 2020;8(21):1408.

44. Libby EK, Pascal KE, Mordechai E, Adelson ME, Trama JP. Atopobium vaginae triggers an innate immune response in an in vitro model of bacterial vaginosis. Microbes and Infection. 2008;10(4):439–46.

45. Borgdorff H, Gautam R, Armstrong SD, Xia D, Ndayisaba GF, van Teijlingen NH, et al. Cervicovaginal microbiome dysbiosis is associated with proteome changes related to alterations of the cervicovaginal mucosal barrier. Mucosal Immunology. 2016;9(3):621–33.

46. Mei L, Wang T, Chen Y, Wei D, Zhang Y, Cui T, et al. Dysbiosis of vaginal microbiota associated with persistent high-risk human papilloma virus infection. Journal of Translational Medicine. 2022;20(1):12.

47. Tabatabaei N, Eren AM, Barreiro LB, Yotova V, Dumaine A, Allard C, et al. Vaginal microbiome in early pregnancy and subsequent risk of spontaneous preterm birth: a case– control study. BJOG: An International Journal of Obstetrics & Gynaecology. 2019;126(3):349–58.

48. Witkin SS, Mendes-Soares H, Linhares IM, Jayaram A, Ledger WJ, Forney LJ, et al. Influence of Vaginal Bacteria and D-and l-Lactic Acid Isomers on Vaginal Extracellular Matrix Metalloproteinase Inducer: Implications for Protection against Upper Genital Tract Infections. mBio. 2013;4(4):e00460–13.

49. Stoyancheva G, Marzotto M, Dellaglio F, Torriani S. Bacteriocin production and gene sequencing analysis from vaginal Lactobacillus strains. Archives of Microbiology. 2014;196(9):645–53.

50. Amabebe E, Anumba DOC. The Vaginal Microenvironment: The Physiologic Role of Lactobacilli. Frontiers in Medicine. 2018;5.

51. Morais IMC, Cordeiro AL, Teixeira GS, Domingues VS, Nardi RMD, Monteiro AS, et al. Biological and physicochemical properties of biosurfactants produced by Lactobacillus jensenii P6A and Lactobacillus gasseri P65. Microbial Cell Factories. 2017;16(1):155.

52. Romero R, Hassan SS, Gajer P, Tarca AL, Fadrosh DW, Nikita L, et al. The composition and stability of the vaginal microbiota of normal pregnant women is different from that of non-pregnant women. Microbiome. 2014;2(1):4.

53. Moscicki A-B, Shi B, Huang H, Barnard E, Li H. Cervical-vaginal microbiome and associated cytokine profiles in a prospective study of HPV 16 acquisition, persistence, and clearance. Front Cell Infect Microbiol. 2020;10(528).

54. Lopes dos Santos Santiago G, Tency I, Verstraelen H, Verhelst R, Trog M, Temmerman M, et al. Longitudinal qPCR Study of the Dynamics of L. crispatus, L. iners, A. vaginae, (Sialidase Positive) G. vaginalis, and P. bivia in the Vagina. PLoS One. 2012;7(9):e45281.

55. Verstraelen H, Verhelst R, Claeys G, De Backer E, Temmerman M, Vaneechoutte M. Longitudinal analysis of the vaginal microflora in pregnancy suggests that L. crispatus promotes the stability of the normal vaginal microflora and that L. gasseri and/or L. iners are more conducive to the occurrence of abnormal vaginal microflora. BMC Microbiology. 2009;9(1):116.

56. Tett A, Pasolli E, Masetti G, Ercolini D, Segata N. Prevotella diversity, niches and interactions with the human host. Nature Reviews Microbiology. 2021;19(9):585–99.

57. van Teijlingen NH, Helgers LC, Zijlstra - Willems EM, van Hamme JL, Ribeiro CMS, Strijbis K, et al. Vaginal dysbiosis associated-bacteria Megasphaera elsdenii and Prevotella timonensis induce immune activation via dendritic cells. Journal of Reproductive Immunology. 2020;138:103085.

58. van Teijlingen NH, Helgers LC, Sarrami-Forooshani R, Zijlstra-Willems EM, van Hamme JL, Segui-Perez C, et al. Vaginal bacterium Prevotella timonensis turns protective Langerhans cells into HIV-1 reservoirs for virus dissemination. The EMBO Journal. 2022;n/a(n/a):e110629.

59. López-Filloy M, Cortez FJ, Gheit T, Cruz y Cruz O, Cruz-Talonia F, Chávez-Torres M, et al. Altered Vaginal Microbiota Composition Correlates With Human Papillomavirus and Mucosal Immune Responses in Women With Symptomatic Cervical Ectopy. Front Cell Infect Microbiol. 2022;12.

60. Audirac-Chalifour A, Torres-Poveda K, Bahena-Román M, Téllez-Sosa J, Martínez-Barnetche J, Cortina-Ceballos B, et al. Cervical Microbiome and Cytokine Profile at Various Stages of Cervical Cancer: A Pilot Study. PLoS One. 2016;11(4):e0153274.

61. France MT, Fu L, Rutt L, Yang H, Humphrys MS, Narina S, et al. Insight into the ecology of vaginal bacteria through integrative analyses of metagenomic and metatranscriptomic data. Genome Biology. 2022;23(1):66.

62. Lebeau A, Bruyere D, Roncarati P, Peixoto P, Hervouet E, Cobraiville G, et al. HPV infection alters vaginal microbiome through down-regulating host mucosal innate peptides used by Lactobacilli as amino acid sources. Nature Communications. 2022;13(1):1076.

63. Chen EY, Tran A, Raho CJ, Birch CM, Crum CP, Hirsch MS. Histological ‘progression’ from low (LSIL) to high (HSIL) squamous intraepithelial lesion is an uncommon event and an indication for quality assurance review. Modern Pathology. 2010;23(8):1045–51.

64. Lopes dos Santos Santiago G, Cools P, Verstraelen H, Trog M, Missine G, Aila NE, et al. Longitudinal Study of the Dynamics of Vaginal Microflora during Two Consecutive Menstrual Cycles. PLoS One. 2011;6(11):e28180.

65. Usyk M, Schlecht NF, Pickering S, Williams L, Sollecito CC, Gradissimo A, et al. molBV reveals immune landscape of bacterial vaginosis and predicts human papillomavirus infection natural history. Nature Communications. 2022;13(1):233.

66. Andralojc KM, Elmelik D, Rasing M, Pater B, Siebers AG, Bekkers R, et al. Targeted RNA next generation sequencing analysis of cervical smears can predict the presence of hrHPV-induced cervical lesions. BMC Med. 2022;20(1):206.

67. Molina MA, Andralojc KM, Leenders WPJ, Huynen MA, Melchers WJG. Cervicovaginal microbial communities and hrHPV infections Sequencing Read Archive: NCBI; 2022 [cited 2022 October 9]. BioProject: PRJNA856437]. Available from: https://www.ncbi.nlm.nih.gov/bioproject/PRJNA856437.

68. Molina MA, Leenders WPJ, Huynen MA, Melchers WJG, Andralojc KM. Longitudinal profiling of the cervicovaginal microbiome Sequencing Read Archive: NCBI; 2022 [cited 2022 October 9]. BioProject: PRJNA888791]. Available from: https://www.ncbi.nlm.nih.gov/bioproject/888791.

