## Supplementary Figure 1 for "Temporal composition of the cervicovaginal microbiome associates with hrHPV infection outcomes in a longitudinal study"

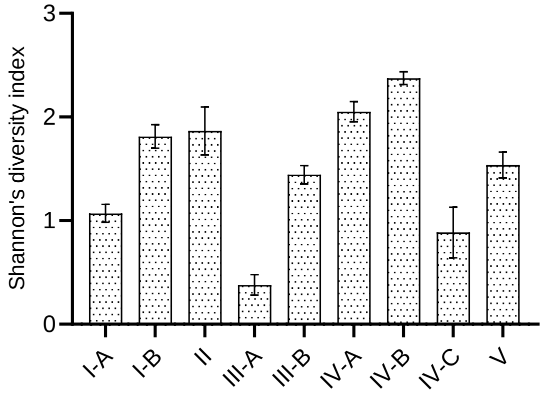


**Supplementary Figure 1. Microbial diversity of microbiomes at V1.**

Analysis of alpha-diversity of the CVM of all participating women at baseline per CSTs as evaluated by Shannon's index (*n* = 141). Error bars represent standard error of the mean ± s.e.m.
