## Supplementary Figure 2 for "Temporal composition of the cervicovaginal microbiome associates with hrHPV infection outcomes in a longitudinal study"

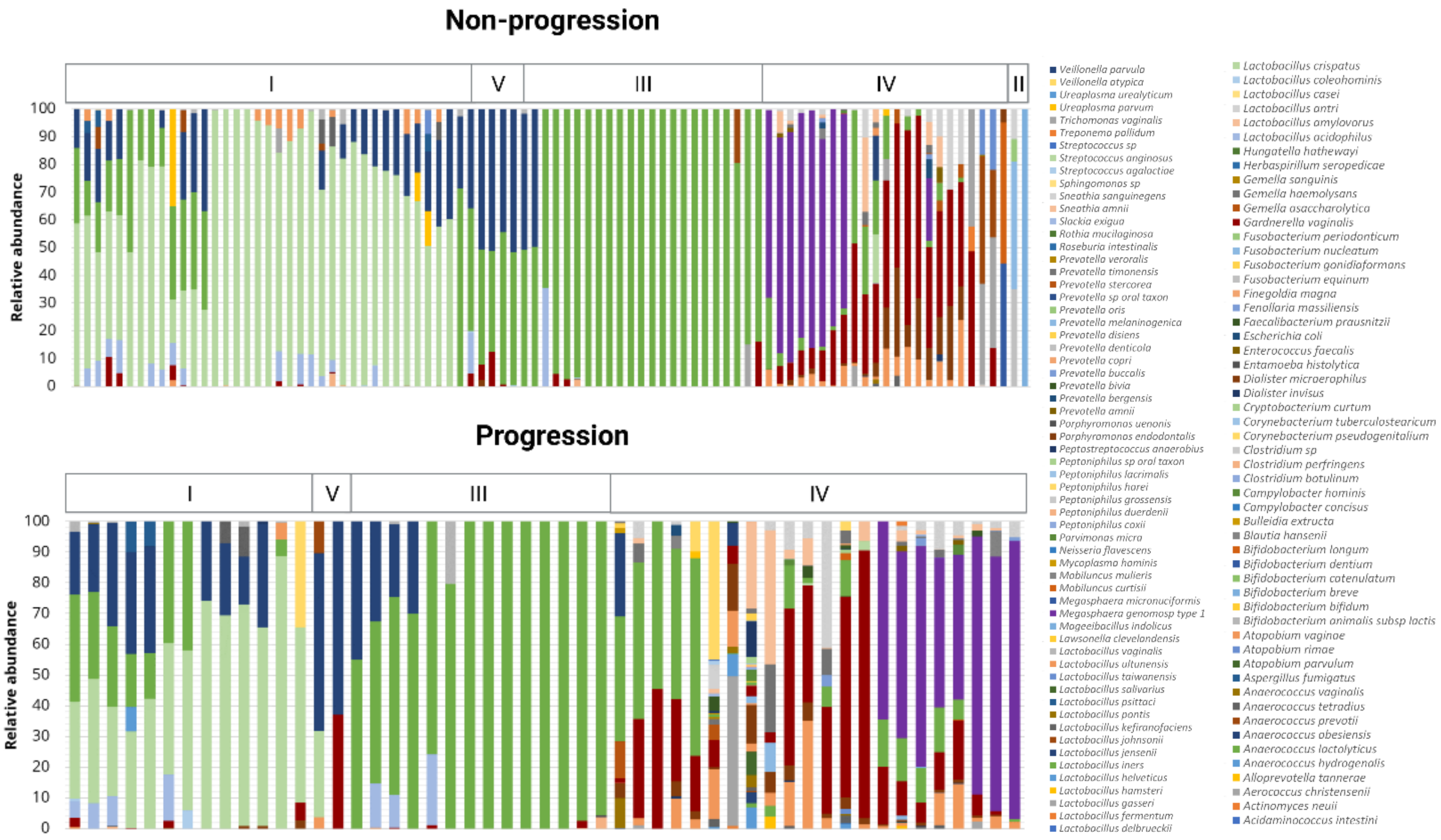


**Supplementary Figure 2. Composition of the microbiomes at V2.**

The cervicovaginal microbiota composition at second collection visit (V2) structures in CSTs based on unsupervised cluster analysis of the microbiomes and is displayed in graph bars (*n* = 141). The NP group is enriched for CSTs I, II, III, and V, while the P group is enriched for CST IV.
