## Supplementary Figure 3 for "Temporal composition of the cervicovaginal microbiome associates with hrHPV infection outcomes in a longitudinal study"

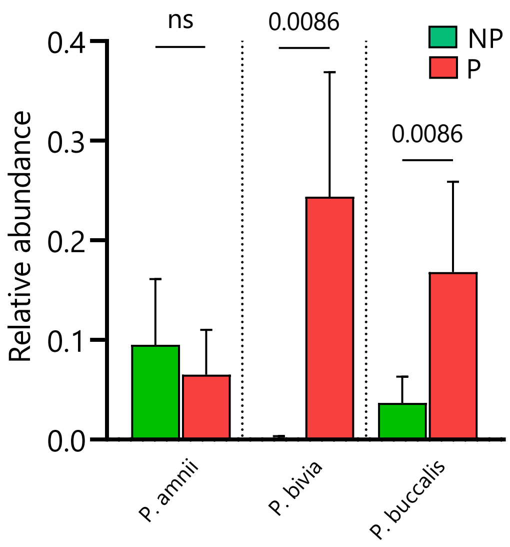


**Supplementary Figure 3. Abundances of *Prevotella* species at V2.**

Analyses of the relative abundances of *Prevotella* species in the CVM of all participating women at second visit (*n* = 141, V2). There is a significantly higher abundance for *P. bivia* and *P. buccalis* in the P group than in the NP group. Differences in relative abundances were analyzed by using a Kruskal-Wallis test followed by the Benjamini-Hochberg test correction for multiple comparisons. *q* values are displayed, and error bars represent standard error of the mean ± s.e.m. NP = non-progression group; P = progression group.
