## Supplementary Figure 4 for "Temporal composition of the cervicovaginal microbiome associates with hrHPV infection outcomes in a longitudinal study"

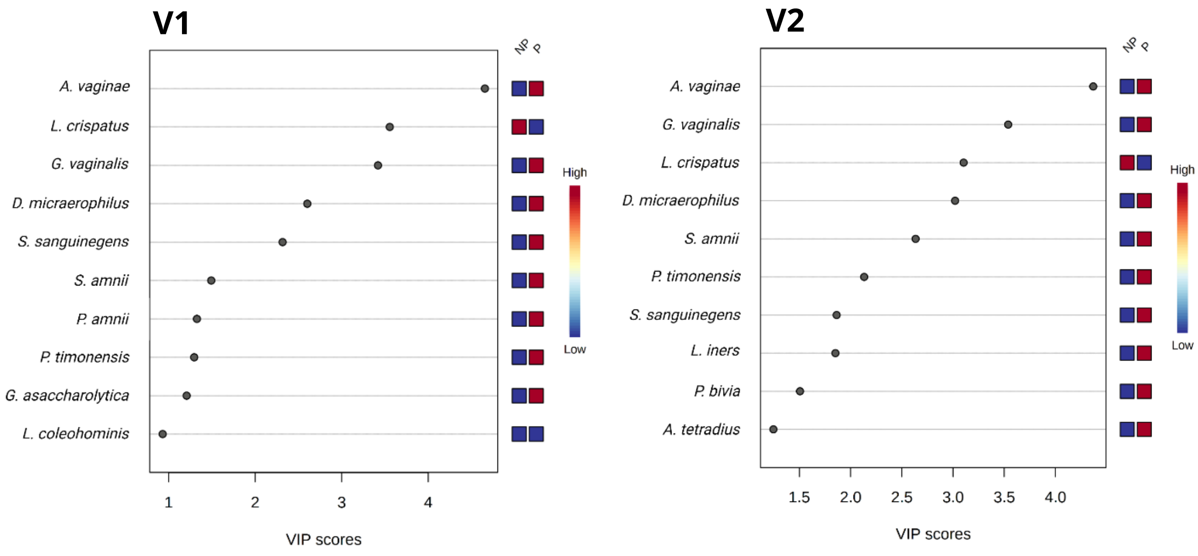


**Supplementary Figure 4. Identification of relevant microbial species in the PLSDA.**

Two component partial least-squares discriminant analysis (PLS-DA) was used to identify the top 10 microbial species that are the most contributory variables in PLSDA C1 for class discrimination between the non-progressive women and progressive women at both collection visits. The index values of the Variable Importance in Projection (VIP) from the PLS-DA component 1 show the species with VIP scores over one. VIP scores are a weighted sum of squares of the PLS loadings. The relative abundance of microbial species is indicated by a colored scale from blue to red representing the low and high, respectively. NP = non-progression group; P = progression group.
